# Anomalous Thermoresponsive Phase Behavior in a Minimal Lattice Model Mimicking Proteins

**DOI:** 10.1101/2025.04.01.646641

**Authors:** Siddhartha Roy, Mantu Santra, Rakesh S. Singh

## Abstract

Proteins are known to phase separate and form biomolecular condensates that play key roles in cellular functions. Therefore, there is now a growing interest in understanding rational design principles for protein sequences that exhibit rich stimulus-responsive phase behavior. Here, we have developed a minimal lattice-gas model and employed Monte-Carlo simulations to capture rich thermoresponsive phase behavior — often observed in computational and experimental studies on disordered proteins. Proteins are modelled as particles with two internal states: a ground state representing the native configuration and a degenerate excited state representing unfolded configurations. The computed phase diagrams reveal an anomalous reentrant phase separation behavior with both upper and lower critical solution temperatures, providing insights into its underlying mechanism. We also explored non-equilibrium effects, such as non-Boltzmann population distribution of native and unfolded states, and enhanced translational diffusion due to non-thermal noise on the phase separation. We find that these factors offer an additional dimension for modulating condensate morphologies. We further extended our model to study phase separation in binary protein mixtures and successfully reproduced complex phase-separated states, including wetted, partially wetted, and segregative phases — observed in computational studies on binary protein mixtures. This model can be easily extended to mimic proteins with multiple internal (partially folded) states, allowing exploration of conformation heterogeneity effects on condensate morphologies. Our findings offer important insights into designing solvent-mediated effective interactions between proteins for controlled phase separation, relevant for engineering functional condensates.

## I. INTRODUCTION

Liquid-liquid phase separation (LLPS) of biopolymers (such as, proteins and RNAs) has been widely recognized as a vital phenomenon responsible for the formation of membraneless organelles — also known as “biomolecular condensates” — in cells^1–3^. These biomolecular condensates are known to play key roles in cellular functions, and therefore, understanding the organization and physical principles underlying (intracellular) LLPS is essential for insights into aging, human health and diseases^4–10^. Furthermore, investigating factors governing LLPS may aid in developing therapeutic strategies to regulate pathological condensates^11^.

Recent experimental and computational studies reveal that LLPS of (bio)polymers can exhibit rich condensate morphologies in response to external environmental factors, such as, temperature (*T*), pressure (*P*), pH, and salt concentrations^12–24^. For example, elevated pressure triggers reentrant phase separation in lysozyme solutions^13^. A similar salt-induced reentrant phase separation has been observed in proteins such as FUS, TDP-43, Brd4, Sox2, and Annexin A11, where phase separation occurs at low and high salt concentrations but is suppressed at intermediate levels due to hydrophobic interactions^25^. Furthermore, proteins (especially intrinsically disordered proteins (IDPs)) exhibit diverse phase separation behaviors in response to temperature^14,26^. These can be broadly classified into three types: (i) upper critical solution temperature (UCST)-type, where proteins phase separate at low temperatures but mix at high temperatures, (ii) lower critical solution temperature (LCST)-type, where proteins remain miscible at low temperatures but demix above the LCST^26^, and (iii) reentrant phase behavior, characterized by UCST at lower temperatures and LCST at higher temperatures, with a miscible region in between^14,27^. For certain proteins and protein-RNA mixtures a reentrant closed-loop phase diagrams featuring LCST at low temperature and UCST at high temperature have also been reported^12,28^. These anomalous thermoresponsive behaviors can be attributed to temperature-induced conformational changes and associated alternation of solvent-mediated biomolecular interactions^14,29^. While UCST-type phase separation is well studied, and sequence determinants are relatively well understood^26,30^, understanding of LCST-type and reentrant phase separations still remains limited^34^.

There is now a growing interest in developing rational design principles for protein sequences that exhibit thermo- or stimulus-responsive (including the non-equilibrium factors, such as non-thermal noise due to the active intracellular processes) phase behavior^17,26,30–33,35–42^. Achieving this requires establishing a direct connection between stimulus-driven changes in a protein’s internal state and the resulting modifications in inter-protein effective interactions^43^. In other words, it will involve a systematic investigation of the following: -the solvent-mediated effective inter-protein interactions that drive the condensate formation and their control via the design of protein sequences and external solvent and thermodynamic factors^44^, (b) the interplay between driven (active) versus spontaneous (passive) conditions required for enabling the formation of rich multiphase condensate architectures^45^, and (c) role of physicochemical diversity arising from multicomponent nature of the intracellular environment on condensate formation. These studies can yield a set of rules regarding homotypic (same molecules) and heterotypic (different molecules) interactions that are likely to be relevant for engineering functional biomolecular condensates^46–50^. Here, we have adopted a top-down approach and constructed a minimal lattice-gas (LG) model for the LLPS of proteins with the aim of understanding the effective interaction (both homotypic and heterotypic) parameter space that would give rise to the above-mentioned diverse phase separation scenarios in response to temperature. Precisely, our aim here is to study the effects of the interplay between parameters that control the single protein thermodynamics (in the absence of inter-protein interactions) and inter-protein effective interactions to the morphology of the biomolecular condensates. As unlike simple molecules, a single protein can exist in many conformational states — the (solvent-mediated) effective interaction between proteins in aqueous environment also depends on their internal conformational states (for example, see Ref. 43). Therefore, protein folding/unfolding thermodynamics at a single protein level, along with the protein’s sequence composition, are known to play key roles in protein’s phase separation^14,17,36,43,51–58^.

In our model, proteins are mimicked via particles with multiple internal states — representing different conformational states. For simplicity, we used a single type of particle, but with two distinct energy states: a non-degenerate ground state (representing native protein conformation) and degenerate excited states (representing entropically-favored unfolded states). The unfolded state has *g*_u_ fold degeneracy and the native protein transitioning to the unfolded state loses *ε* energy. The parameters *g*_u_ and *ε* dictate the single protein thermodynamics and the unfolded-state population can be given as, 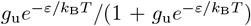) (*k*_B_ is the Boltzmann constant), in the absence of inter-protein interactions. Of course, the distribution of the folded and unfolded conformations at a single protein level could get significantly altered in a condensed phase due to the inter-protein interactions. In addition, we have explicitly accounted for the protein’s internal state dependent inter-protein interactions and studied the effect of heterotypic (same type of molecules but in different conformational states) interaction strength, temperature, and unfolded state degeneracy *g*_u_ on the phase behaviour of the system.

This study reveals an anomalous reentrant LLPS with both UCST and LCST, shedding light on its underlying mechanisms. We have further investigated non-equilibrium effects, such as deviations from Boltzmann population distributions and enhanced translational diffusion driven by non-thermal noise, demonstrating their role in shaping LLPS morphologies. Finally, we extended our model to binary protein mixtures and captured diverse phase-separated states — wetted, partially wetted, and segregative phases — consistent with recent computational studies^35,59^. These insights may pave the way for designing protein sequences in conjunction with external environmental factors (such as solvent and thermodynamic conditions) for tunable phase behavior, essential for intracellular condensate regulation.

The organization of the rest of the paper is as follows. In Section II we discuss our lattice model and the Monte-Carlo methods to simulate this model system at both equilibrium and driven out-of-equilibrium conditions. In Section III A we present the unfolded-state degeneracy induced criticality and phase transition in the protein’s conformational space. The LLPS for different heterotypic interaction parameters and temperature conditions is discussed in Sections III B and III C. In Section III D we present the phase separation scenario in a binary protein mixture, and the major conclusions from this work are summarized in Section IV.

## II. MODEL AND METHOD DETAILS

### A. Generalized Lattice-Gas Model

In our minimal LG model proteins are represented by particles with multiple internal states — non-degenerate ground state which represents the folded (or, native) protein configuration and degenerate higher energy states representing unfolded protein configurations. A schematic representation of the same with two internal states, native and unfolded, is shown in Fig. 1. This system is represented by a square lattice with *N* sites and each site is assigned to a variables (occupation number) *n*_*i*_ which takes values 1 or 0 depending on the presence or the absence of a particle on site *i*, respectively. The internal states of particles are represented by a variable *σ*_*i*_, which takes on the *p* discrete values 1, 2,…, *p*, where *p* is the total number of protein’s internal (conformational) states. As is evident, *σ*_*i*_ is only defined when *n*_*i*_ is 1 (i.e. site is occupied with a particle). For the sake of simplicity, in this work we have considered only two protein internal states (*p* = 2) — native and unfolded (denoted by *n* and *u*, respectively). Of course, proteins in intermediate states with partially folded and unfolded domains can easily be modelled by adding intermediate states between the native and the unfolded states — like a Pott’s model. We can now write down the Hamiltonian of the system as:

**FIG. 1.**
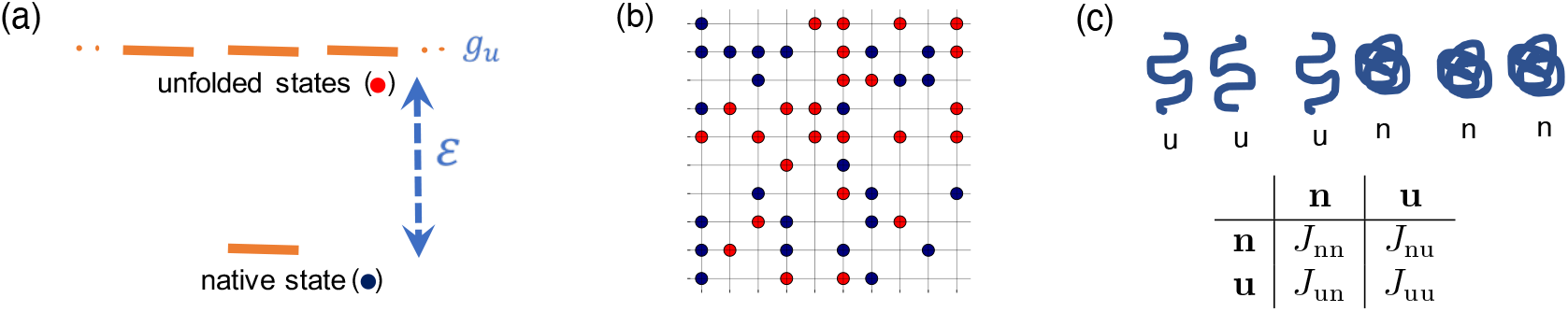
(a) The proteins are represented with particles having two internal states, native (n, denoted by blue filled circle) and unfolded (u, denoted by red filled circle). The degeneracies of the native and unfolded states are 1 and *g*_*u*_, respectively. *ε* is the internal interaction energy difference between the native and unfolded states that account for the loss of internal weak interactions on protein unfolding. (b) A schematic representation of the lattice system with randomly dispersed native and unfolded proteins. The empty sites represent the solvent. (c) Internal conformational state-dependent interactions between proteins (top) and the corresponding interaction matrix representing the strength of the unfolded-unfolded (*J*_uu_), native-unfolded (*J*_nu_), and native-native (*J*_nn_) interactions (botttom).

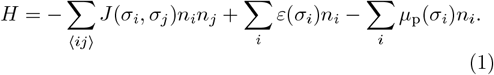

Here, *J (σ*_*i*_, σ_*j*_) represents the protein’s internal state dependent interaction between the *i*^th^ and its nearest neighbor *j*^th^ site, and for two internal state proteins we have three possibilities — *J*_uu_, *J*_nn_ and *J*_un_ — denoting unfolded-unfolded, native-native and unfolded-native interactions, respectively (interaction matrix is a symmetric 2 *×* 2 matrix), and ⟨*ij*⟩ denotes nearest neighbour pairs. *ε*(*σ*_*i*_) (with σ_*i*_ = n or u) is the internal energy of the native and unfolded states (see Fig. 1b). We have set ε_*n*_ = 0 (the native state energy) and *ε*_*u*_ = −2.0*J*_uu_ (unfolded state energy with respective to the native state). The values of the chemical potential at the *i*^th^ site, *µ*_p_(*σ*_*i*_), is set to 2.0*J*_uu_ (this value ensures an one-to-one mapping of the mean-field behavior of the LG and Ising systems with only one energy scale *J*_uu_).

In this model, we have multiple energy scales and their relative importance to the bulk phase behvaior can be controlled by changing *T* or *g*_*u*_. The heterotypic interaction *J*_un_ is varied during the study and homotypic interactions, *J*_uu_ and *J*_nn_, are set to 1.0 and 0.5, respectively, throughout this study. Here, homotypic and heterotypic interactions refer to intermolecular interactions between the proteins in the same versus different conformational states, respectively. The relative values of *J*_uu_ and *J*_nn_ are judiciously chosen based on the fact that in the native (folded) state many of the protein’s interaction sites are either occupied internally or buried inside the core, and therefore, not available for inter-protein interactions resulting a weaker inter-protein interactions. However, it is worth noting that the relative strength of *J*_uu_ and *J*_nn_ depends on protein sequence details and on external (solvent and thermodynamic) conditions, and may display a diverse range. Although, these diverse possibilities, once known computationally or experimentally, can be easily incorporated in our model, in the subsequent sections we have discussed results only for above-mentioned homotypic interaction parameters. Similar type of minimal models have previously been used to explore phase separation and pattern formation under both equilibrium and non-equilibrium conditions, though in contexts distinct from those examined in this work^60–65^.

### B. Computational Method Details

We employed Monte-Carlo (MC) simulations in canonical ensemble to sample the configuration space. As is evident from the model, we needed to incorporate two types of moves — folding-unfolding conformational change (for sampling conformational space) and translation — of particles on lattice sites. As we are interested in the phase separation phenomena, the fraction of proteins (or, the occupied sites, denoted as *ϕ*_p_) is a conserved variable. However, the fraction of proteins in the unfolded or folded (*ϕ*_n_ and *ϕ*_u_ states, respectively) is a non-conserved variable, and can vary with time or MC steps. We have used Kawasaki exchange dynamics^66^ for the translational move of the occupied and unoccupied lattice sites. In Kawasaki dynamics, the positions of a pair of occupied-unoccupied sites or two occupied sites with different (native or unfolded) internal states are exchanged, so that the particle concentration is locally conserved. For the non-equilibrium cases (such as deviations from Boltzmann unfolded/native population distribution and enhanced translational diffusion driven by non-thermal noise), however, only the positions of a pair of occupied-unoccupied sites, not two occupied sites with different internal states, are exchanged. The exchange rate depends only on the energy difference (Δ*E*) involved in this process as the proposed move acceptance probability (*p*_acc_) is given as, *p*_acc_ = min[1, exp(−*β*Δ*E*)]; *β* = 1*/k*_B_*T*. Here, the chemical potential plays a role analogous to the external field. We have used a simple Metropolis criterion^67^ for the folded-unfolded transition moves. The system is equilibrated for at least 5*×*10^6^ MC steps with the equal frequency of the translational and conformational moves. We applied the periodic boundary condition along both (*x* and *y*) directions. We chose a rectangular simulation box with lattice sizes *L*_*x*_ = 200 and *L*_*y*_ = 50 to stabilize the planar interface between the coexisting phases along *x*-direction^68^. We observed the formation of droplets with roughly circular interfaces in a square box at lower protein concentrations. In the subsequent sections, temperature in reported in units of *J*_uu_*/k*_B_, that is, reduced temperature (*T* ^*^) is defined as *k*_B_*T/J*_uu_.

## III. RESULTS AND DISCUSSION

### A. Unfolded state degeneracy-driven phase transition in the conformational composition space

As the inter-protein (effective) interactions are coupled to their internal conformational states, we first explored the control of the protein system’s (global) conformational ensemble through single protein’s unfolded state degeneracy, *g*_u_ — a single protein property — at a given temperature. Here, we fixed the protein concentration ⟨*ϕ*_p_⟩ to 0.5 along with the inter-protein homotypic interactions (*J*_nn_ and *J*_uu_), and varied *g*_u_ for different heterotypic interactions *J*_un_, probing the dependence of the average fraction of proteins in the unfolded state (⟨ *ϕ*_u_ ⟩) on *g*_u_ at a fixed *T* (see Fig. 2). We note that at lower temperatures and for weaker heterotypic interactions (characterized by a smaller *J*_un_ value), ⟨ *ϕ*_u_ ⟩ exhibits a discontinuous change with increasing *g*_u_ (Fig. 2a)). Additionally, we observe a hysteresis in *ϕ*_u_ when the control parameter *g*_u_ is reversed for *T* ^*^ = 0.3 (see Fig. S1a in the Supplementary Material) which indicates a first-order-like native-to-unfolded population transition. Furthermore, at higher temperatures, for a given low *J*_un_ value, the folded-to-unfolded transition is more gradual with no associated hysteresis on reversal (see Fig. 2a and Fig. S1b in the Supplementary Material), resembling a transition at supercritical conditions. At an intermediate temperature, we observe a critical-like scenario in the conformational composition space, where the slope of the order parameter ⟨*ϕ*_u_⟩ with respect to the control parameter *g*_u_ diverges. On increasing the heterotypic interaction strength *J*_un_ we observe that the native-unfolded population transition gradually softens and changes in a continuous manner in the *T* -range studied with a relatively sharper change at lower *T* s compared to the higher (see Figs. 2b and 2c).

**FIG. 2.**
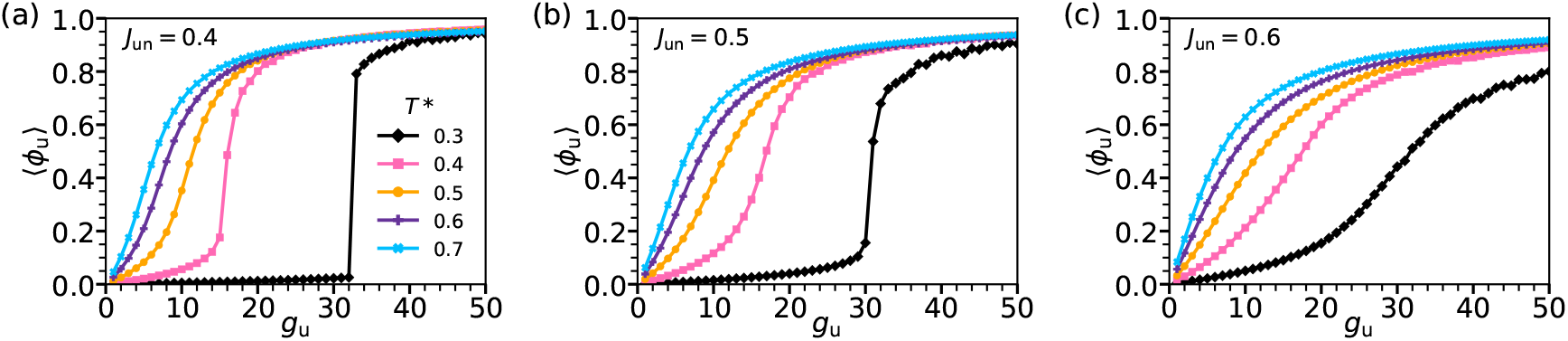
The dependence of the fraction of unfolded state proteins (⟨*ϕ*_u_⟩) in the system on the degeneracy of the unfolded state *g*_u_ at different reduced temperatures (*T* ^*^) for the heterotypic interaction parameter *J*_un_ = 0.4 (a), 0.5 (b), and 0.6 (c). Here, *ϕ* _p_ is set to 0.5. We note a discontinuous (first-order-like) transition between the native and unfolded states at lower *T* ^*^ and *J*_un_, and rather a continuous transition at higher *T* ^*^ and *J*_un_.

We further computed the order parameter susceptibility 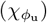, defined as, 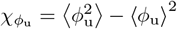, to understand the nature of the underlying free energy surface [*F* (⟨*ϕ*_u_) vs. ⟨*ϕ*_u_⟩] and its dependence on the parameter *g*_u_ (see Fig. S2 in the Supplementary Material). As expected from the equations of state shown in Fig. 2a, 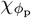 exhibits a maximum near the transition *g*_u_, which becomes more pronounced with increasing system size, suggesting a possible divergence in the thermodynamic limit. The above results suggest that the *g*_u_-driven folded-to-unfolded transition (and *vice-versa*) bears a close resemblance to the van der Waals equation of state and may belong to the Ising universality class. However, we have not performed a detailed finite-size scaling analysis or calculated the critical exponents to definitively establish the universality class as this is beyond the scope of the present work.

### B. Liquid-liquid phase separation

#### 1. Reentrant phase separation

After gaining an understanding of the unfolded-state degeneracy-dependent system’s conformational behavior, we have elaborately explored here the phase behavior under monotonic variation of temperature for various inter-protein heterotypic interactions *J*_un_ and different unfolded-state degeneracies. In Fig. 3a, we present a *J*_un_ − *T* ^*^ phase diagram, illustrating the different regions (phase-separated and well-mixed) for *g*_u_ = 20. It is evident from the figure that our model exhibits reentrant phase behavior. Starting with a completely phase-separated state P1 at low *T* where proteins are predominantly (*>* 50%) in the native state, an increase in *T* (while keeping other interaction parameters fixed) drives the system to transition into a mixed state P2, where the proteins are homogeneously dispersed. Interestingly, upon further increasing *T*, the system transitions again to a phase-separated state P3, where the proteins predominantly exist in the unfolded state. As expected, at sufficiently high temperatures, guided by the strength of inter-particle interactions, the system eventually becomes homogeneous (named as P4). The snapshots of configurations at state points belonging to P1, P2, P3 and P4 phases, along with the respective composition profiles, are shown in Figs. 3b and 3c for *J*_un_ = 0.2 and 0.6, respectively. We conducted our studies for various initial conditions, such as all proteins being in the native state, in the unfolded state, and randomly distributed and found that the phase diagram remains nearly invariant on changing the initial configuration. This phase behavior shows a close resemblance with the reported reentrant phase separation on IDPs and other (bio)polymers with both UCST and LCST^12,14^. Here, we observe two UCST lines — named as UCST-I and UCST-II — between the P1-P2 and P3-P4 phases, respectively, in addition to the LCST line between the P2 and P3.

**FIG. 3.**
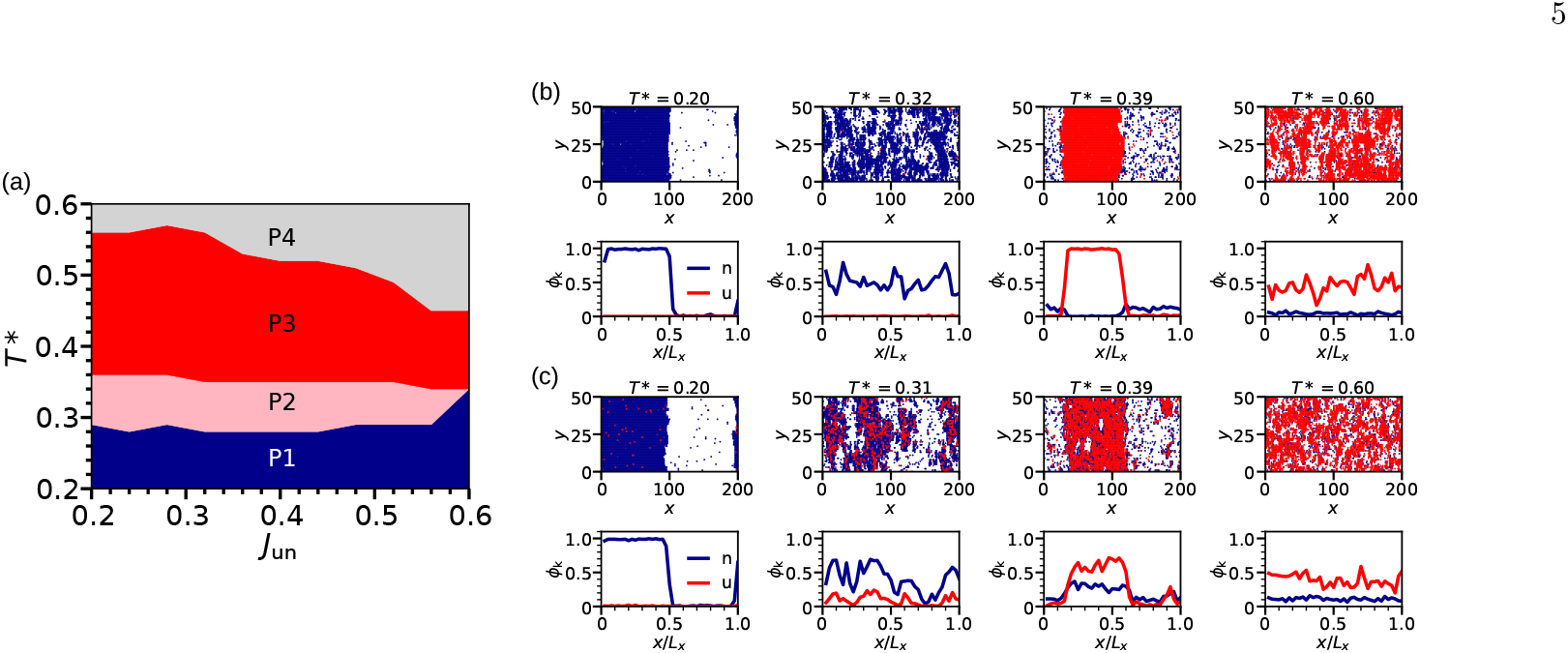
(a) Phase diagram showing the different (phase separated and mixed) states in the *T* ^*^ − *J*_un_ plane for *g*_u_ = 20 and *ϕ*_p_ = 0.5. Note, the reentrant phase behavior where the phase separated phase P1, where the proteins are predominantly in the native state, undergoes a mixing transition to phase P2 on increasing *T* and this homogeneous state with mixed folded and unfolded protein conformations again gets de-mixed (P3) on further increasing the temperature. P3 phase predominantly contains unfolded state proteins. P4 represents a homogeneous dispersed phase where proteins are predominantly in the unfolded state. The snapshots, along with the corresponding density profiles, at representative state points belonging to P1, P2, P3 and P4 phases are depicted in (b) and (c) for *J*_un_ = 0.2 and 0.6, respectively. In the snapshots, the blue and red colors represent the native and unfolded proteins, respectively.

It is evident from Figs. 3b and 3c that, although the phase separation scenario qualitatively remains the same (i.e., reentrant), the native/unfolded composition differs significantly with the change in heterotypic interaction *J*_un_ for a given choice of homotypic interactions and temperature (see also Fig. S3 in the Supplementary Material). For instance, for *J*_un_ = 0.2, the P2 phase predominantly contains native proteins, whereas in the P3 phase, the phase-separated region consists solely of unfolded proteins, with the folded proteins dispersed in the homogeneous medium — indicative of a segregative phase separation (see Fig. 3b (top panel); *T* ^*^ = 0.39). This behavior is also evident from the corresponding density profiles of native and unfolded conformations shown in Fig. 3b (bottom panel). In contrast, for *J*_un_ = 0.6, the P2 and P3 phases exhibit a mixed (native/unfolded) composition — representing an associative phase separation (see Fig. 3c; *T* ^*^ = 0.39 and the corresponding density profile shown in the bottom panel of this figure)^12,69,70^.

We have further explored the effects of *g*_u_ on phase separation morphologies by computing the phase diagram for a higher *g*_u_ value (*g*_u_ = 30; see Fig. 4), where the unfolded state is more stable (lower free energy) compared to the previous case (*g*_u_ = 20). We find that as *g*_u_ increases, the reentrant mechanism gets suppressed, with the intermediate P2 phase region shrinking and eventually disappearing at higher *J*_un_ values. In Figs. 4b and 4c, we present the representative snapshots along with the corresponding density profiles for *J*_un_ = 0.2 and *J*_un_ = 0.6, respectively. At lower *J*_un_ values, the *T* -dependent reentrant phase separation mechanism resembles that observed for the previously studied case (*g*_u_ = 20). However, at higher *J*_un_ values (*J*_un_ *>* 0.45), the native-dominated P1 phase continuously evolves into the unfolded-dominated P3 state without encountering any intervening homogeneous phase (Fig. 4c). In this case, phase boundaries are determined based on the dominance of one conformational state over the other. For example, if *ϕ*_u_ *<* 0.5, the phase-separated region is marked in blue, indicating predominantly native conformations, whereas if *ϕ*_u_ *>* 0.5, it is marked in red, indicating unfolded-state dominance.

**FIG. 4.**
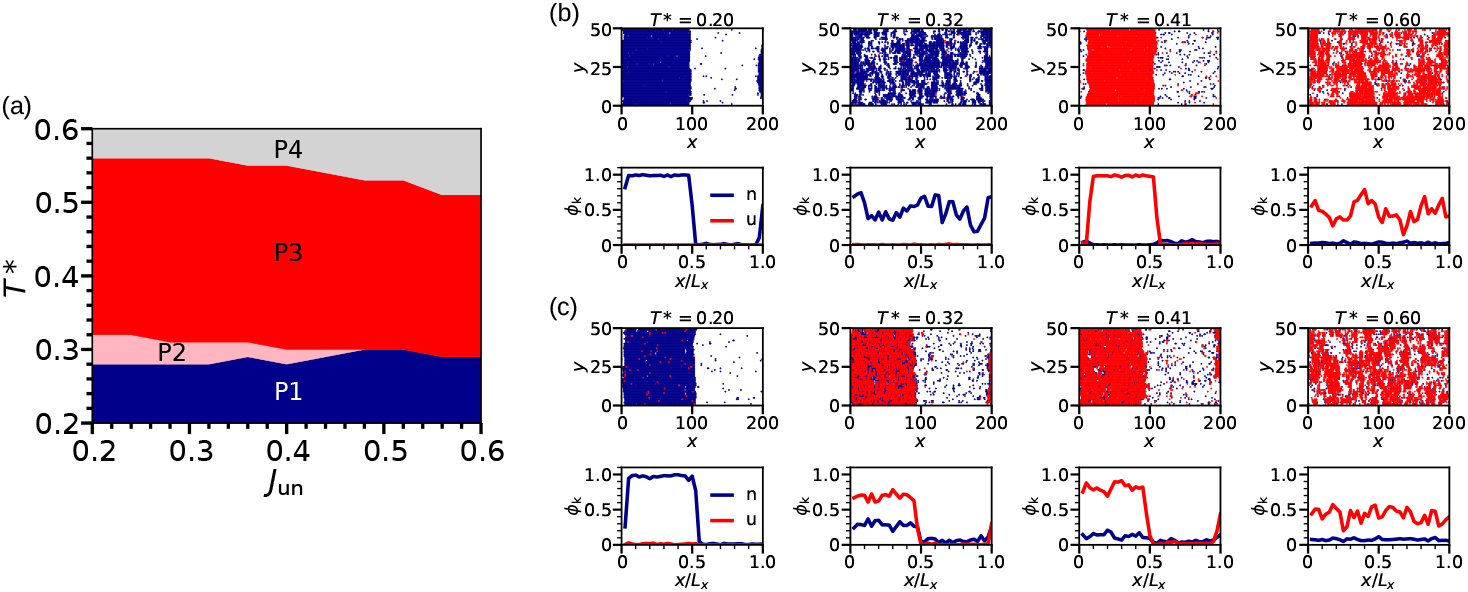
(a) Phase diagram showing the different (phase-separated and mixed) states in the *T* ^*^ − *J*_un_ plane for *g*_u_ = 30 and *ϕ*_p_ = 0.5. The representative snapshots of different phases, along with the corresponding density profiles of phases are depicted in (b) and (c) for *J*_un_ = 0.2 and 0.6, respectively. In the snapshots, the blue and red colors represent the native and unfolded proteins, respectively.

These results provide a reliable protocol to control the segregative versus associative LLPS scenarios either by tuning the effective interaction between proteins (or between different protein types in the case of a protein mixture) or by altering single-protein unfolding-folding thermodynamics via *g*_u_ and *ε* parameters.

#### 2. Origin of the reentrant phase behavior

To gain deeper insights into the observed reentrant behavior and its origin rooted in conformational and interaction diversity, we investigated the *T* dependence of ⟨*ϕ*_u_⟩, and average energy per particle (⟨*e*_p_ ⟩) for different values of *J*_un_ (see Fig. 5). As *T* increases, the system transitions from native to unfolded protein conformational states (Fig. 5a). This transition becomes continuous at high *J*_un_. A similar trend is observed in the temperature variation of ⟨*e*_p_⟩ (Fig. 5b). We note that the de-mixed to mixed transition of the native protein phase-separated state does not lead to significant changes in either the protein’s conformational state or the system’s energy. However, at the reentrant (mixed to demixed) transition temperature, *ϕ*_u_ changes abruptly, resulting in a sudden increase in the system’s interaction energy (especially for lower *J*_un_ values). This clearly suggests that the reentrant transition is entropy-driven. For higher *J*_un_ values, however, the energy (like ⟨*ϕ*_u_⟩) increases gradually due to the progressive change in the protein conformational state, making the intervening native-rich mixed phase region narrower and eventually disappears for *J*_un_ *>* 0.6.

**FIG. 5.**
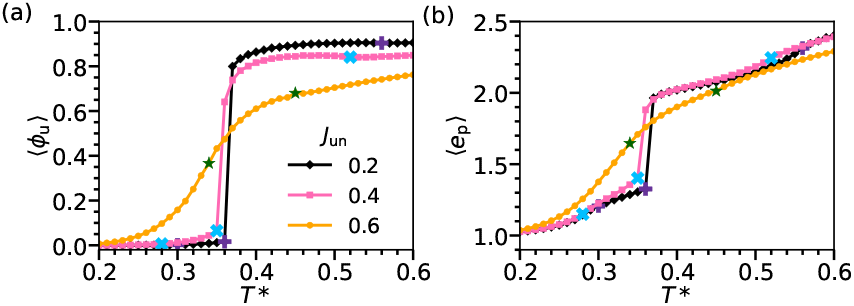
Temperature dependence of the fraction of proteins in the unfolded state, ⟨*ϕ*_u_ ⟩ (a) and interaction energy per protein *e*_p_ (b) for the heterotypic interaction *J*_un_ = 0.2, 0.4, and 0.6. The unfolded state degeneracy *g*_u_ is set to 20. In (a) and (b), the different symbols mark the (mixed/de-mixed) transition temperatures in the phase diagram shown in Fig. 3a for the respective *J*_un_ values.

The above observations suggest that the reentrant behavior arises due to the presence of multiple energy scales and the *T* -dependence of one energy scale over the other. This can be understood by invoking the (inter-particle) interaction strength-dependent critical point as the critical temperature is proportional to the coupling parameter strength. At lower temperatures, all proteins predominantly exist in the energetically favored native state and a weaker inter-(native) protein interaction (*J*_nn_ *< J*_uu_) leads to a lower critical temperature (denoted as *T*_cn_). As *T* increases, the system becomes supercritical with respect to *T*_cn_, where the proteins undergo a mixing transition (see Fig. S4 in the Supplementary Material). Upon further increasing *T*, the proteins transition (either gradually or discontinuously) into the unfolded-dominated state (see Fig. 5a) where the inter-protein interactions are much stronger compared to the native-dominated state. This transition alters the critical temperature either gradually or abruptly (depending on *J*_un_), shifting it to a higher temperature (denoted as *T*_cu_). Consequently, the system becomes sub-critical again but this time with respect to *T*_cu_, resulting in reentrant phase behavior — particularly at lower or intermediate *J*_un_ values. For higher *J*_un_ values, the reentrant behavior vanishes because the native-to-unfolded transition becomes gradual, leading to a continuous increase in the dominance of stronger inter-protein interactions, and in turn, the critical temperature. As a result, the system remains subcritical until it reaches the *T*_cu_ for the given *J*_un_.

### C. Phase separation at driven out-of-equilibrium conditions

In living cells, active processes — such as ATP-driven molecular motors and cytoskeletal dynamics — can introduce non-thermal noise influencing phase separation. Furthermore, crowding and other factors associated with the cytosol medium (such as interaction of biopolymers with crowders) can also give rise to a sluggish translational diffusion and may also contribute to the non-equilibrium proteins conformational population, and in turn to the LLPS phenomena^71–73^. In this section, we have explored two non-equilibrium scenarios: (a) the protein conformational states follow non-Boltzmann population distribution, and (b) translation diffusion of proteins is enhanced compared to the thermal equilibrium case due to the presence of non-thermal noise.

#### 1. Protein conformational states follow non-Boltzmann population distribution

To create a non-equilibrium conformational state distribution, an external biasing potential (*E*_b_) is incorporated into the Boltzmann factor of the Metropolis acceptance criterion as *p*_acc_ = min[1, exp[−*β*(Δ*E* − *E*_b_)]]; Δ*E* is the difference in the interaction energy associated with the MC move involving protein’s conformational change (see Section II B). For a positive *E*_b_ the population is biased towards the unfolded state and vice-versa (see Fig. S5a in the Supplementary Material). In Fig. 6a we show the phase diagram under this non-equilibrium conditions where the conformational states follow non-Boltzmann distribution. We observe that, for the case of unfolded state population bias, reentrant behavior disappears in the way that P1 transitions to P3 on increasing *T* without the intervening mixed P2 phase, resulting in the disappearance of both the UCST-I and LCST. For low *J*_un_ values, this (P1 to P3) transition is abrupt, however, for higher values, this transition is rather gradual (as discussed in Section III B 1, in the case of gradual transition (for higher *J*_un_ values), phase boundaries are marked on the basis of the dominance of one conformational state over the other). For the native-biased case (*E*_b_ *<* 0), however, we find that the range of the intervening mixed phase region gets expanded. This expansion mainly results from the shift of the LCST line to higher temperature (the UCST-I line remains almost unchanged). In addition, the P3 state disappears for higher *J*_un_ values (Fig. 6a(ii)), and therefore, for the native-biased scenario only at low *J*_un_ values, reentrant LLPS is observed. We also note that the phase boundary separating P3 and P4 move towards higher *T* on biasing the conformation population towards the native state (Fig. 6a(ii)).

**FIG. 6.**
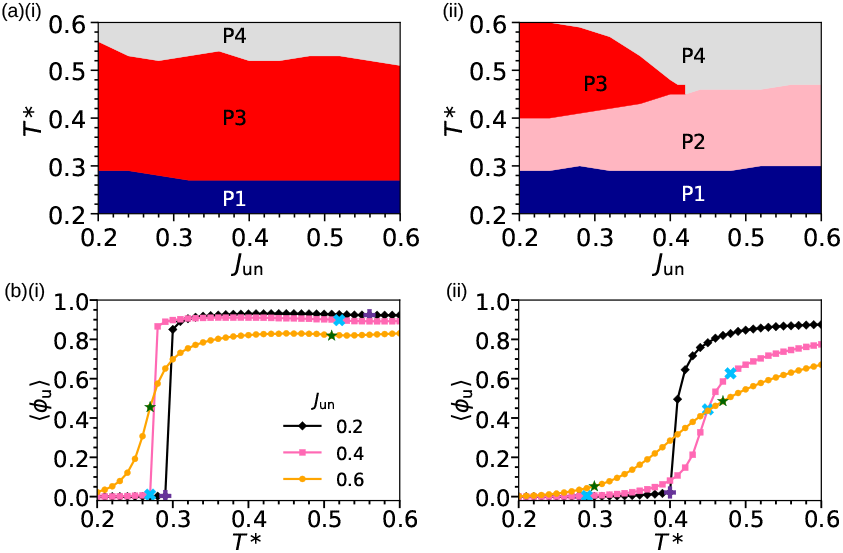
(a) The phase diagram in the *T* ^*^ − *J*_un_ plane depicts the various phases from P1 to P4 for the biasing potential *E*_b_ = 0.2 (i) and 0.2 (ii) [see Fig. 3 for representative structures of P1, P2, P3 and P4 phases]. As described in the main text, a positive biasing potential favors the higher energy unfolded state and vice-versa. We note that the phase separation scenario is strongly sensitive to *E*_b_, or the strength of the external driving force. (b) The temperature dependence of the unfolded state population ⟨*ϕ*_u_ ⟩ for different *J*_un_ values for *E*_b_ = 0.2 (i) and −0.2 (ii). The phase boundary between the homogeneous P2 and P4 phases indicates the dominance of the native protein in P2 and unfolded in P4.

To understand the local structural origin of these non-equilibrium factor-driven observed alterations in the phase behavior, in Fig. 6b we show the *T* -dependent ⟨*ϕ*_u_⟩ for both native- and unfolded-bias scenarios. In the unfolded-bias case (Fig. 6b(i)), ⟨*ϕ*_u_⟩ rises sharply (and abruptly for lower *J*_un_ values) and this transition occurs at lower temperatures compared to the equilibrium and native-bias cases. This increase of the (strongly-interacting) unfolded-state proteins gives rise to an increase of the (effective) critical temperature, resulting the system to remain subcritical over a wide temperature range compared to the native-bias case. In contrast, in the native-bias case, the growth of ⟨*ϕ*_u_⟩ with *T* is suppressed, requiring higher temperatures for comparable population increase. As a result, once system becomes supercritical with respect to *T*_cn_ it remains supercritical across a broader temperature range compared to the thermal equilibrium and unfolded-bias scenarios. We also note that the increase of ⟨*ϕ*_u_⟩ with *T* is sharper, for a given *J*_un_, for the unfolded-population bias case (Fig. 6b(ii)) than the native-bias. The absence of the reentrant phase behavior at higher *J*_un_ for the native-bias case could be attributed to this gradual and relatively weak *T* -dependence of ⟨*ϕ*_u_⟩, which does not allow the dominance of the (strong) interaction-driven structuring of unfolded proteins over the destabilizing thermal effects even at relatively lower temperatures. In other words, the critical temperature increases gradually due to the increase of the strongly interacting unfolded protein population with *T*, however, the system always remains supercritical with respective to the respective critical point due to the concurrent increase of thermal fluctuations.

#### 2. Translational mobility is enhanced due to the non-thermal noise

We further explored the effects of translational diffusion enhancement on phase separation behavior. One possible cause of this enhancement in intracellular enhancement is the presence of non-thermal noise, in addition to thermal fluctuations. To model this non-equilibrium scenario, we introduce an external biasing potential *E*_b_ into the Boltzmann factor of the Metropolis acceptance criterion, modifying the Kawasaki exchange dynamics in the similar way as discussed above (Section III C 1). A positive biasing energy enhances translational mobility of proteins. Interestingly, we observed that, along with the translation diffusion change, the biasing potential also alters the relative unfolded/native population (see Fig. S5b in the Supplementary Material). In Fig. 7a we present the phase diagram for this non-equilibrium scenario. We observe that the native phase separated state P1 almost disappears, despite no change in the native-state population. We attribute this disappearance of P1 to the fact that the enhanced translation diffusivity puts the system effectively at a higher temperature (*T > T*_cn_, for this given value of *E*_b_) than the actual *T* and hence no phase separation. Therefore, interestingly, at low *J*_un_ and sufficiently large *E*_b_, the system may exhibit a reverse reentrant behavior: it is mixed at low *T*, transitions to a de-mixed state upon heating (due to the dominance of the stronger inter-protein interactions), and then gets mixed again at a higher *T*. This behavior shows a close resemblance with the closed loop phase separation with LCST at low *T* and UCST at high *T*^12,14^.

**FIG. 7.**
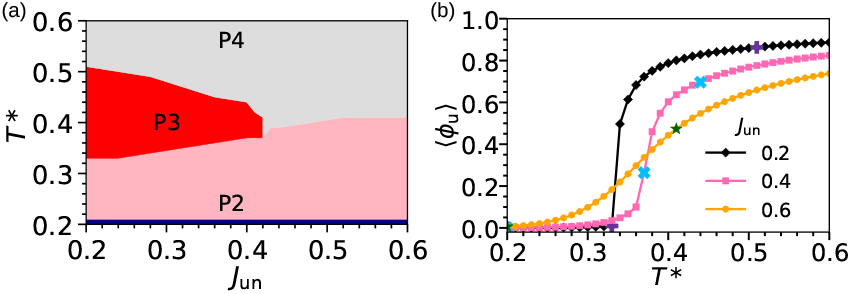
(a) The phase diagram in the *T* ^*^ − *J*_un_ plane depicts the various phases from P1 to P4 for the biasing potential *E*_b_ = 0.2 which enhances the diffusivity of proteins in the system. It is evident from the figure that the phase separation scenario is strongly sensitive to *E*_b_. (b) The temperature dependence of the unfolded state population ⟨*ϕ*_u_⟩ for different *J*_un_ values.

For larger *J*_un_ values, the system remains mixed across the entire temperature range probed. The reason for this can again be attributed to the softening of the *T* - dependence of *ϕ*_u_ for higher *J*_un_ (see Fig. 7b) — as discussed in the previous section. The above observations suggest that the interplay between temperature, non-equilibrium conditions, and heterotypic interaction strength offers a robust means of controlling phase morphology and mixture composition. Impact of macro-molecular crowding on protein-protein interactions, and in turn, on LLPS has not been explored in this work, and could be an interesting avenue for future research^72^.

### D. Phase separation in binary protein mixture

We have now investigated a binary mixture system consisting of two different types of proteins. Recent studies by Chew et al.^35^ on binary protein mixtures — where proteins were evolved in their amino acid sequence — reveal a rich phase separation behavior, including selective wetting of one component by the other, depending on their interactions. Their findings indicate that larger differences in homotypic interactions favor de-mixing, with the inner phase of the condensate always composed of the protein species exhibiting stronger homotypic interactions. In our model, this binary mixture system (comprising, say, proteins A and B) can be mimicked by restricting folding-unfolding transitions, making the particles non-interconvertible. Consequently, the lattice dynamics involve only translational motion via Kawasaki exchange dynamics. This model provides an effective framework to study phase behavior in scenarios where (major) conformational transitions are inaccessible due to insufficient thermal energy during the phase separation process. Here, we have set *g*_u_ = 1 and *E* = 0 in the model Hamiltonian (Eq. 1). The fractions of both A and B proteins (*ϕ*_A_ and *ϕ*_B_, respectively) are set to 0.25 so that the total protein fraction *ϕ*_p_ (= *ϕ*_A_ + *ϕ*_B_) is 0.5.

Figure 8 presents the phase diagram in the *T* − *J*_AB_ plane for the binary protein mixture system. As in the previous case, we set here also the homotypic interactions to *J*_AA_ = 1.0 and *J*_BB_ = 0.5, varying the heterotypic interaction *J*_AB_. This yields a diverse range of phase behavior, including partially wetted, fully wetted, and segregative (along with the mixed or homogeneous) phases (see representative snapshots in Fig. 8c and corresponding density profiles in Fig. 8d). To construct the phase diagram, we generated an initial configuration in a phase-separated state with randomly mixed A and B proteins (see Fig. 8b), equilibrated it at the desired temperature, and analyzed the final structure. At low-*T* and low *J*_AB_ we observe an interesting partially wetted phase where one side of the B-rich regions is in contact with the A-rich region, while the other side is in contact with the solvent (or vacuum in our model). Increasing *J*_AB_ at low *T* leads to a fully wetted phase (Fig. 8c), and further increasing *J*_AB_ (≥ 0.6) results in the dispersion of wetting-phase B proteins into the solvent — giving rise to a segregative phase where A proteins are phase *T* * separated in a homogeneous mixture of B proteins and solvent. For low *J*_AB_, the segregative phase emerges at higher temperatures (see Fig. 8). These phase-separated morphologies closely resemble those reported by Chew et al.^35^, except for the partially wetted phase. Wetting of one phase by another reduces the surface free energy and is known to play an important role in facilitating phase transitions^74–76^.

**FIG. 8.**
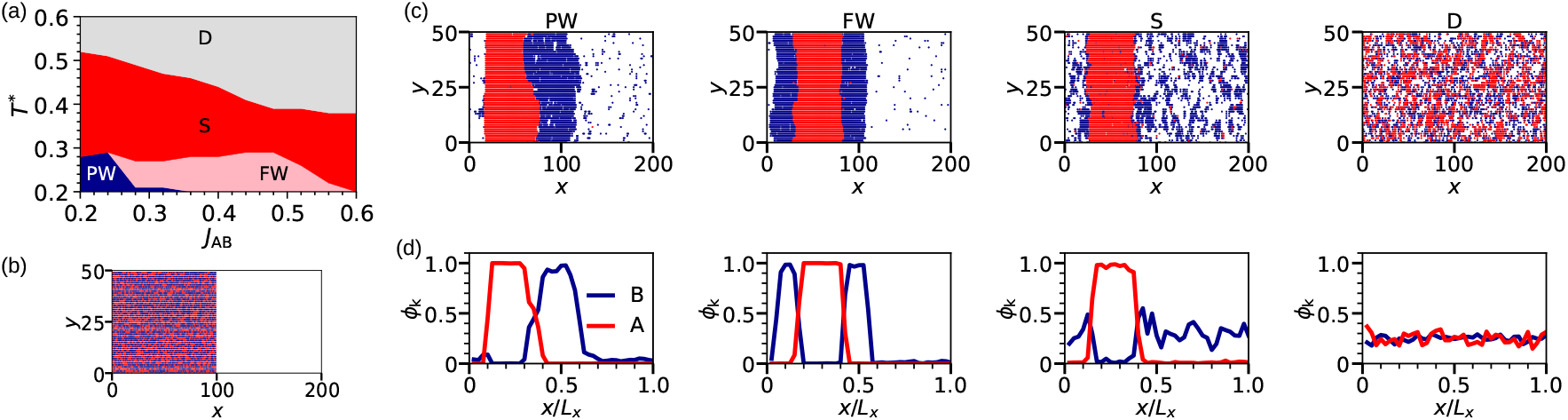
(a) The phase diagram in the *T* ^*^ − *J*_*AB*_ (*T* ^*^ ≡ *k*_B_*T/J*_AA_) plane showing the partially-wetted (PW), fully-wetted (FW), segregative (S), and mixed or fully dispersed (D) phases. In the segregative phase, strongly-interacting A proteins are phase separated and B proteins are dispersed inside the medium. (b) Initial configuration with a randomly mixed folded-unfolded protein composition is used for computing the phase diagram. The snapshots of configurations representing different phases and the respective A-B composition profiles are shown in (c) and (d), respectively. In the snapshots, the blue and red color represent A and B type proteins, respectively. We note that, at low temperatures, the PW phase transitions to FW on increasing *J*_*AB*_, and on further increasing *J*_*AB*_ the FW phase undergoes a transition to the S phase.

In Fig. 9 we present the variation of interaction energy per protein, *e*_p_, with temperature for different heterotypic interaction strengths, *J*_AB_. The clear crossover in the *T* -dependence of *e*_p_ near the transition temperatures (characterized by the sharp change in the slope of the *T* -dependence of *e*_p_; marked with different symbols for different *J*_AB_s in Fig. 9) indicates that signatures of entropy-driven phase transitions are encoded in the variation of energy of the system with *T*. The crossovers are more pronounced for lower *J*_AB_. We also examined the dependence of the phase diagram on initial conditions by considering different starting phase-separated states — partially wetted, fully wetted, and reverse-wetted phases (see Fig. S6 in the Supplementary Material for details). Our results show that the low-temperature region of the phase diagram is particularly sensitive to initial conditions, and in addition to the partially and fully wetted phases, a reverse-wetted phase is also observed to exist, likely as a metastable state, at low and intermediate *J*_un_. This finding suggests a potential existence of exotic metastable phase-separated states in real systems with distinct cellular functionalities.

**FIG. 9.**
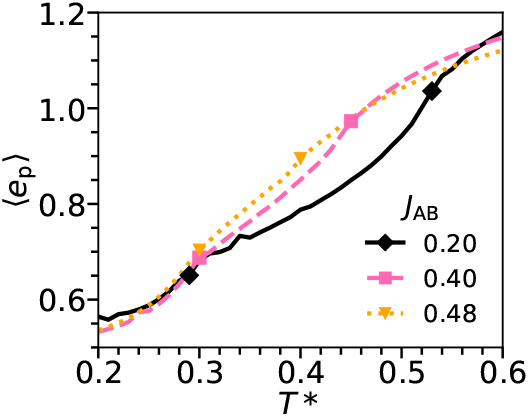
(a) The variation of the average energy per protein (⟨*e*_p_⟩) with *T* ^*^ for different heterotypic interaction parameter (*J*_AB_) values. The diamond, square and triangular symbols indicate the transition temperatures for *J*_AB_ = 0.20, 0.40 and 0.48, respectively. The homotypic interaction parameters are set to *J*_AA_ = 1.0 and *J*_BB_ = 0.5.

The above results suggest that the interplay between thermodynamic conditions and the effective interactions between constituents can give rise to a diverse range of phase-separated morphologies in a multicomponent system. Interaction heterogeneities is intrinsic to a multi-component intracellular environment, and *P, T*, pH and salt-induced conformational fluctuations and transitions may also add to this complexity.

## IV. CONCLUSIONS

In this work, we have used a minimal statistical mechanical model based on the LG Hamiltonian to capture the diverse phase-separated architectures of biomolecular condensates — including reentrant LLPS in response to temperature — observed in experimental and computational studies of coarse-grained protein models. Proteins are represented as particles with two internal states: a non-degenerate ground state corresponding to the folded (native) protein configuration and a degenerate excited state representing unfolded configurations. The parameters in our model can be categorized into two types: (i) those governing single-protein thermodynamics, which control the population distribution between folded and unfolded states, and (ii) inter-protein interaction parameters. The resulting phase diagram, mapped in the temperature-interaction parameter space, reveals an anomalous reentrant LLPS featuring both UCST and LCST. Our analysis provides key insights into the underlying mechanisms driving this behavior. Additionally, we identify (effective) interaction parameters that lead to associative and segregative phase separation.

Given that the intracellular environment operates out of equilibrium, we further investigated non-equilibrium effects — such as non-Boltzmann population distributions of native and unfolded states and enhanced translational diffusion due to non-thermal noise — on LLPS. We find that these non-equilibrium factors provide an additional dimension for modulating condensate morphologies. Moreover, we extended our model to study LLPS in binary protein systems by restricting internal conformational transitions and allowing only translational dynamics. The model successfully reproduces complex phase-separated morphologies, such as, wetted, partially-wetted, and segregative phases — recently reported in computational studies of binary coarse-grained protein systems^35,59^.

It is worth noting here that our model can be readily extended to proteins with multiple internal states (e.g., partially folded or disordered domains — a common feature among phase separating proteins, such as IDPs) by incorporating additional degrees of freedom in a *q* state Potts-like model^77,78^. This extension would enable a more detailed exploration of the role of structural disorder and folded domains in biomolecular condensate formation and morphology. Furthermore, environmental factors such as pH or salt concentration can be incorporated by modulating the relative stability of native and unfolded states along with the inter-particle interactions. While solvent effects are implicitly included in our model, a more realistic approach would involve explicit solvent interactions within the two-state LG Hamiltonian, a direction currently under investigation. Finally, our study provides a foundation for guiding future coarse-grained computational and experimental efforts to design protein sequences that optimize both single-protein thermodynamics (folding-unfolding equilibrium) and inter-protein interactions (homotypic and heterotypic) to achieve targeted self-assembled phase-separated architectures. This ability to control biomolecular condensate morphology is crucial for the functional regulation of biomolecular condensates.

## ACKNOWLEDGMENTS

R.S.S. acknowledges financial support from DST-SERB (Grant No. CRG/2023/002975). S.R. acknowledges financial support from IISER Tirupati. M. S. acknowledges financial support from DST-SERB (Grant No. SRG/2020/001385). The computations were performed at the IISER Tirupati computing facility.

## Supplementary Material

**FIG. S1.**
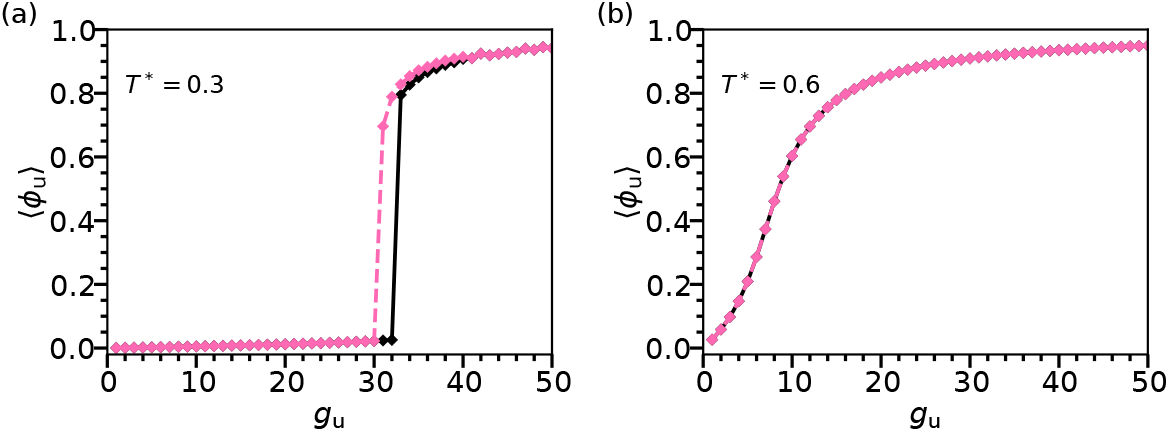
The dependence of the average fraction of proteins in the unfolded state ⟨*ϕ*_u_⟩ on the unfolded state degeneracy *g*_u_ at *T* ^*^ = 0.3 (a) and *T* ^*^ = 0.6 (b). The black line represents the forward path (increasing *g*_u_) and the dashed pink line represents the path on reversal. We note the presence of hysteresis in the *g*_u_ dependence of ⟨*ϕ*_u_⟩ for *T* ^*^ = 0.3 and its absence for *T* ^*^ = 0.6. Here the homotypic interaction parameters are set to *J*_nn_ = 0.5 and *J*_uu_ = 1.0 and the heterotypic interaction *J*_un_ is 0.4.

**FIG. S2.**
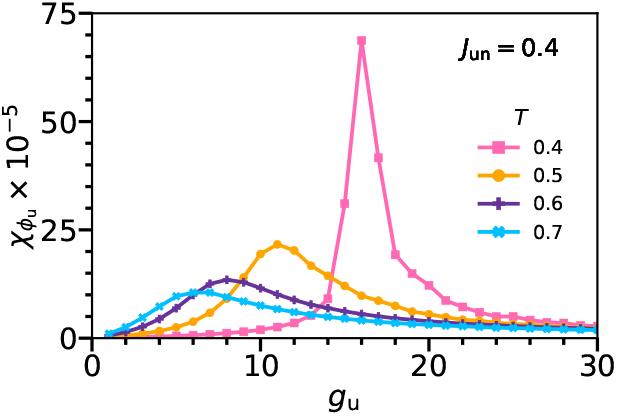
The order parameter (fraction of proteins in the unfolded state) susceptibility 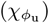 defined as, 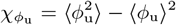, as a function of the unfolded-state degeneracy *g*_u_ at different temperatures. We note a sharp increase of 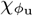 on approaching the temperature where *ϕ*_u_ changes sharply with *g*_u_ (see Fig. 2a in the main text).

**FIG. S3.**
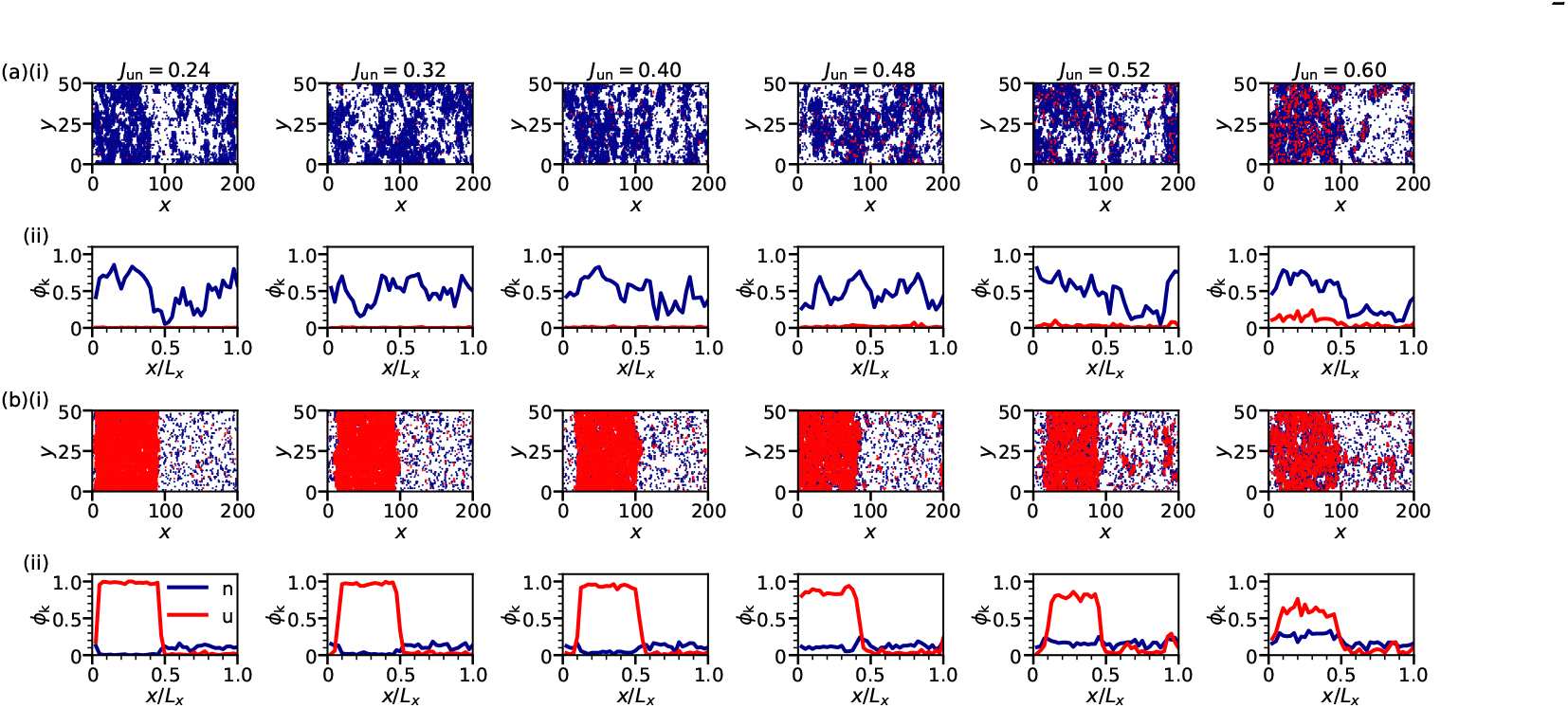
The snapshots (i) and corresponding composition profiles (ii) of the system containing native and unfolded proteins in P2 (a) and P3 (b) phases for *g*_u_ = 20 and *ϕ*_p_ = 0.5 (see Fig. 3 in the main text). Here, at a given temperature (*T* ^*^ = 0.3 for P2, and 0.4 for P3), we have varied *J*_un_ to probe the sensitivity of the system’s composition on the heterotypic interaction parameter strength.

**FIG. S4.**
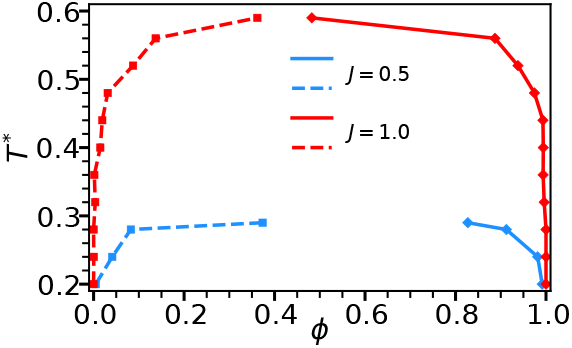
The phase diagram in temperature-composition (*T* ^*^ − *ϕ*_p_) plane for a lattice-gas system where all the proteins are in the unfolded state (i.e. *J*_uu_ = 1.0; indicated by red color), and the native state (*J*_nn_ = 0.5; indicated by blue color).

**FIG. S5.**
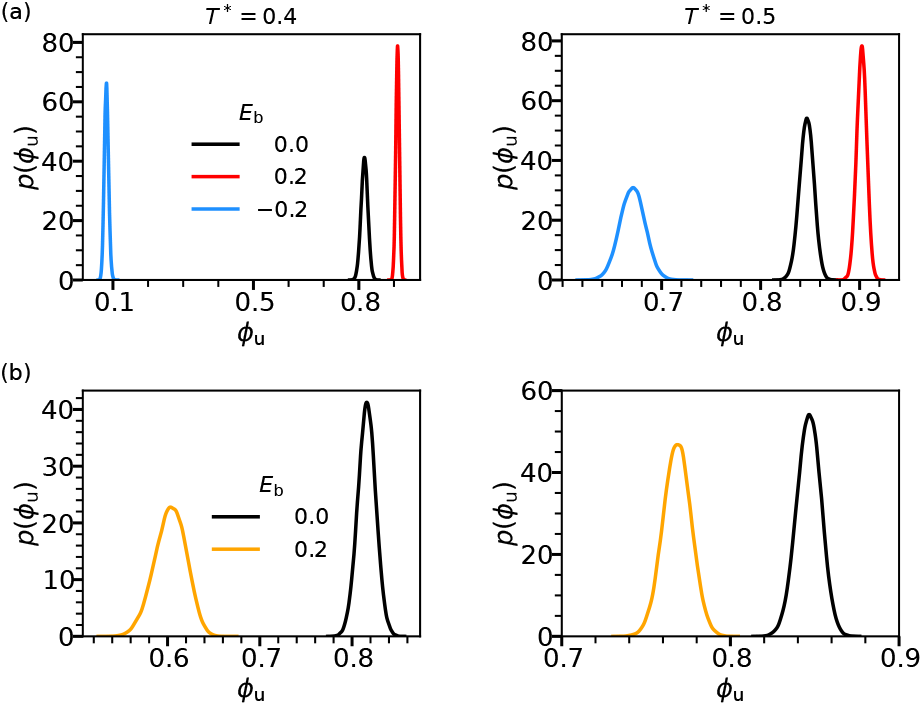
The sensitivity of the unfolded-state population distribution *ϕ*_u_ on the biasing potential *E*_b_ for conformational (a) and translation (b) bias cases for two temperatures are shown. As expected, for the conformational-bias case, the unfolded-state population depends on both *E*_b_ and *T*. Interestingly, however, for the translational bias case, the biasing energy *E*_b_ alters the translation mobility as well as the unfolded-state population. Here, *J*_un_ is set to 0.4.

**FIG. S6.**
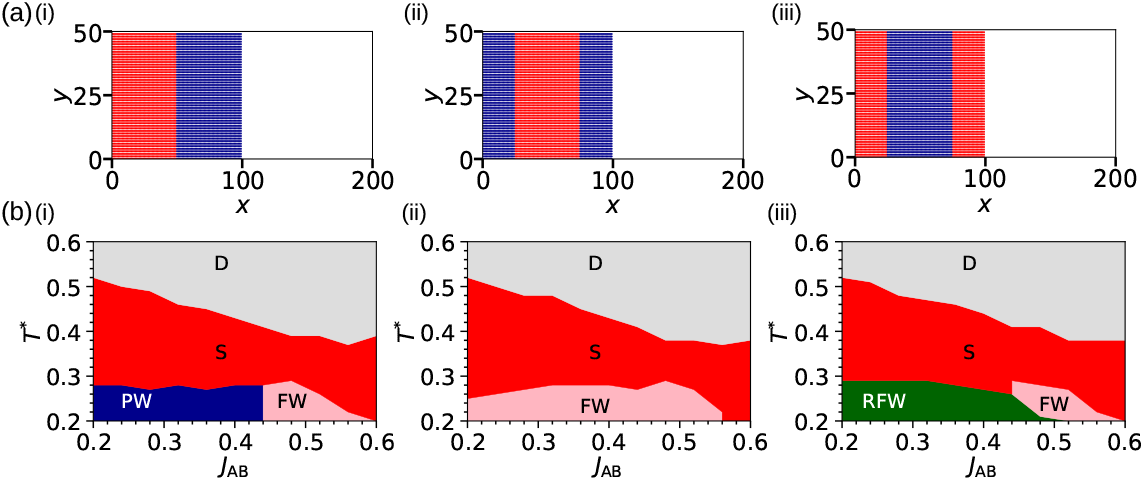
The dependence of the phase diagram on the initial condition is shown for the binary protein system. (a) We have considered three different initial conditions: (i) partially-wetted (PW), (ii) fully-wetted (FW), and (iii) reverse-wetted (RW). In the case of reverse wetting scenario, the weakly interacting B proteins are wetted by the strongly interacting A proteins. (b) The corresponding phase diagrams in *T* ^*^ − *J*_AB_ are shown, We note the sensitivity of the phase diagram on the initial condition, especially in the low *T* region of the phase diagram.

